# Rapid systemic responses of Arabidopsis to waterlogging stress

**DOI:** 10.1101/2023.03.22.533859

**Authors:** María Ángeles Peláez-Vico, Adama Tukuli, Pallav Singh, David G Mendoza-Cózatl, Trupti Joshi, Ron Mittler

## Abstract

Waterlogging stress (WLS) negatively impacts the growth and yield of crops resulting in heavy losses to agricultural production. Previous studies revealed that WLS induces a systemic response in shoots that is partially dependent on the plant hormones ethylene and abscisic acid. However, the role of rapid cell-to-cell signaling pathways, such as the reactive oxygen species (ROS) and calcium waves, in systemic responses of plants to WLS is unknown at present. Here we reveal that an abrupt WLS treatment of *Arabidopsis thaliana* plants growing in peat moss triggers systemic ROS and calcium wave responses, and that the WLS-triggered ROS wave response of Arabidopsis is dependent on the ROS generating RESPIRATORY BURST OXIDASE HOMOLOG D (RBOHD), calcium-permeable channels GLUTAMATE-LIKE RECEPTOR 3.3 and 3.6 (GLR3.3 and GLR3.6), and aquaporin PLASMA MEMBRANE INTRINSIC PROTEIN 2;1 (PIP2;1) proteins. We further show that WLS is accompanied by a rapid systemic transcriptomic response that is evident as early as 10 min following waterlogging initiation, includes many hypoxia-response transcripts, and is partially dependent on RBOHD. Interestingly, the abrupt WLS of Arabidopsis resulted in the triggering of a rapid hydraulic wave response and the transient opening of stomata on leaves. Taken together, our findings reveal that the initiation of WLS in plants is accompanied by rapid systemic physiological and transcriptomic responses that involve the ROS, calcium, and hydraulic waves. These findings reveal that systemic plant responses to WLS are rapid and at least partially dependent on cell-to-cell signaling mechanisms.

## INTRODUCTION

As our climate changes, the frequency and intensity of weather episodes, such as floods and heavy downpours, gradually increase (Bailey-Serres et al., 2019; Masson-Delmotte et al., 2021). Floods and heavy downpours can cover entire fields causing complete or partial submergence of crops, and/or create lasting conditions of waterlogging stress (WLS) by soaking the soil with water for extended periods of time (Voesenek and Bailey-Serres, 2015; Loreti et al., 2016; Pucciariello and Perata, 2017; Sasidharan et al., 2018, 2021). These conditions limit oxygen availability to the root system (waterlogging), or the entire plant (submergence), induce hypoxia- and/or anoxia-response mechanisms, and negatively impact crop growth and yield, resulting in heavy losses to agricultural production (Bailey-Serres et al., 2019).

Waterlogging can rapidly occur under field conditions following a sudden downpour, or as a result of advancing flood water, and create a situation in which the roots are subjected to hypoxia, while the shoots are not (Voesenek and Bailey-Serres, 2015). Previous work has shown that waterlogging causes local hypoxia-driven responses in the roots, and systemic responses in the shoots that involve adjustments in carbohydrate metabolism, ubiquitin-dependent protein degradation, hormonal responses, and many other molecular and metabolic responses (Hsu et al., 2011). In addition, some of the systemic responses induced by WLS were found to be altered in mutants deficient in ethylene and abscisic acid (ABA) signaling (Hsu et al., 2011; Tsai et al., 2014).

Among the first responses to anoxia conditions in roots or shoots of plants are the inhibition of mitochondrial respiration, the activation of calcium signaling, and the accumulation of reactive oxygen species (ROS; Voesenek and Bailey-Serres, 2015; Loreti et al., 2016; Pucciariello and Perata, 2017; Sasidharan et al., 2018, 2021; Yang et al., 2022). Recent studies demonstrated that the vacuolar H^+^/calcium transporter <u>CA</u>TION/PROTON E<u>X</u>CHANGER 1 (CAX1) and the RESPIRATORY BURST OXIDASE HOMOLOGs D and F (RBOHD and RBOHF) play important roles in these responses and that changes in calcium signaling and ROS are important for triggering different anoxia response mechanisms, including transcript accumulation, and in some instances aerenchyma formation (Liu et al., 2017; Yang et al., 2022). The function of RBOHD, RBOHF, and CAX1 was also shown to be required for plant acclimation to anoxia stress (Liu et al., 2017; Yang et al., 2022).

As WLS can occur rapidly in the field (*e.g.,* Voesenek and Bailey-Serres, 2015), cause rapid calcium- and RBOH-driven ROS production in roots (Liu et al., 2017; Yang et al., 2022), and trigger systemic responses in the shoot (Hsu et al., 2011), we hypothesized that WLS could trigger a rapid systemic signaling response that involves the ROS wave. The ROS wave is a cell-to-cell signaling mechanism that depends on RBOHD and RBOHF function and propagates through the vascular bundles and/or mesophyll cells of plants from the site of its stimulation (by abiotic or biotic stress) to the entire plant within minutes (Zandalinas et al., 2020a; Zandalinas et al., 2020b; Fichman and Mittler, 2021; Fichman et al., 2022; Mittler et al., 2022). Integrated with the calcium and electric waves, the ROS wave is also required for the activation of many molecular, physiological, and metabolic responses of systemic tissues, as well as the overall acclimation of plants to different stresses (*e.g.,* Kollist et al., 2019; Fichman et al., 2020a).

Here we show that an abrupt WLS treatment of *Arabidopsis thaliana* plants growing in peat moss triggers the systemic ROS and calcium wave responses, and that the WLS-triggered ROS wave response of Arabidopsis is dependent on RBOHD, the calcium-permeable channels GLUTAMATE-LIKE RECEPTOR 3.3 and 3.6 (GLR3.3GLR3.6), and the aquaporin/peroxiporin PLASMA MEMBRANE INTRINSIC PROTEIN 2;1 (PIP2;1). We further show that WLS is accompanied by a rapid systemic transcriptomic response that is partially dependent on RBOHD. Interestingly, the abrupt WLS of Arabidopsis resulted in the triggering of a rapid hydraulic wave response and the transient opening of stomata on leaves. Taken together, our findings reveal that WLS is accompanied by rapid systemic molecular and physiological responses that involve the ROS, calcium, and hydraulic waves, and that the ROS wave triggered upon waterlogging stress in Arabidopsis is dependent on RBOHD, GLR3.3GLR3.6 and PIP2;1 function. These findings suggest that systemic plant responses to WLS are rapid and at least partially dependent on cell-to-cell signaling.

## RESULTS

### Inducing waterlogging stress in Arabidopsis

To mimic conditions that accompany a sudden event of WLS, caused by a heavy downpour or advancing flood water, we grew Arabidopsis plants in peat soil under controlled growth conditions, allowed the water content of the peat soil to reach 54 ± 2% of full water capacity, and subjected plants to WLS by rapidly watering plants until the water level reached all the way to the top of the peat soil (rapidly equilibrate to about 100% water capacity; Figure 1A). This treatment resulted a significant decrease in oxygen levels measured with an oxygen electrode, as early as 10 min following WLS initiation (Figure 1B). The experimental system used in this study resulted therefore in a state in which the top part of the plant remained in air, while to entire root system of the plant experienced hypoxia stress caused by the WLS treatment (Figure 1).

**Figure 1.**
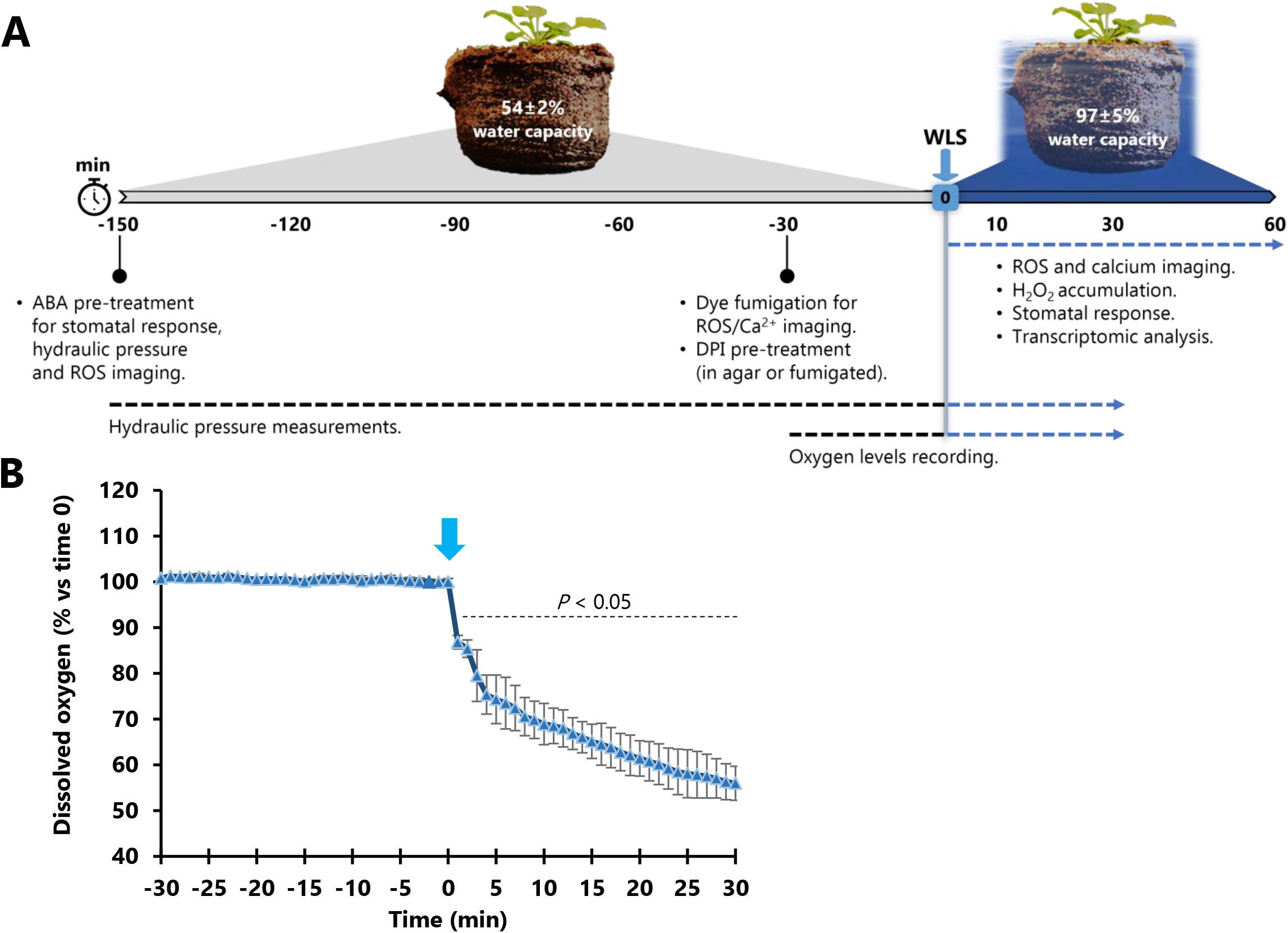
Experimental design and measurements of oxygen levels. (A) The experimental design used to induce waterlogging stress (WLS) in Arabidopsis. Plants grown in peat soil at a defined water content were watered until water reached the top of the peat soil level and analyzed for systemic responses as described in the text. Gray and blue sections indicate pre- and post-WLS respectively; blue arrow indicates WLS treatment application (0 min). (B) Measurements of oxygen level at the middle of the peat soil prior to and during the WLS treatment (applied at 0 min). Statistical analysis was performed with a two-sided Student’s t test (\**P* < 0.05; n = 3). Abbreviations: ABA, abscisic acid; CT, control; DPI, diphenyleneiodonium; H_2_O_2_, hydrogen peroxide; ROS, reactive oxygen species; WLS, waterlogging stress.

### Waterlogging stress triggers the systemic ROS and calcium wave responses

Using the experimental system shown in Figure 1 we studied whether WLS triggers the ROS and calcium waves in the systemic tissues of plants subjected to WLS. As shown in Figure 2A and 2B, WLS resulted in the activation of systemic ROS (Figure 2A) and calcium (Figure 2B) wave responses that were detected in the upper parts of plants within 10-(calcium) and 20-(ROS) min post WLS application, respectively (measured using our live whole-plant imaging method; Fichman et al., 2019). To examine whether the ROS wave response detected in plants subjected to WLS resulted in enhanced accumulation of H_2_O_2_, which plays a key role in regulating plant responses to stress (Mittler et al., 2022), we also measured H_2_O_2_ levels in shoots of plants subjected to WLS using the Amplex Red method (Fichman et al., 2022). As shown in Figure 2C, H_2_O_2_ levels were transiently elevated in shoots of plants subjected to WLS for 30 min and declined at 60 min post stress application.

**Figure 2.**
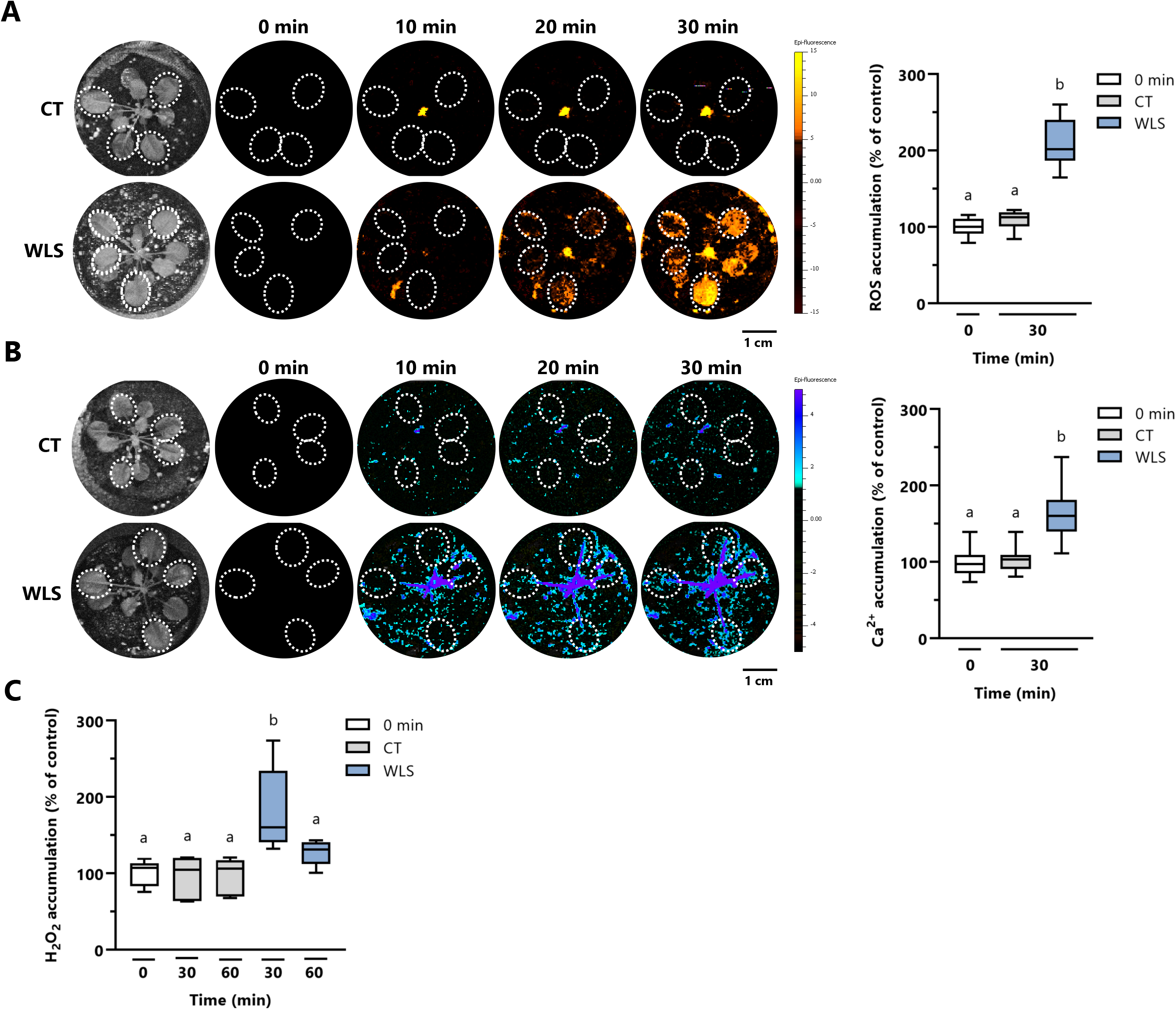
Imaging of systemic ROS and calcium accumulation in wild type plants subjected to an abrupt waterlogging stress (WLS) treatment. (A) *Arabidopsis thaliana* plants were fumigated with H_2_DCFDA and imaged for systemic ROS accumulation in response to WLS. (B) Same as in (A), but for plants fumigated with Fluo-4-AM and imaged for systemic calcium accumulation. Representative time-lapse images of whole plant ROS or calcium accumulation in treated and untreated Arabidopsis plants are shown alongside bar graphs of combined data from all plants used for the analysis at the 0- and 30-min time points in (A) and (B). Regions of interest are indicated by white ovals in the images. Magnification bars in (A) and (B) are 1 cm. All experiments were repeated at least three times with eight plants per repeat. Data is shown as box and whisker plots with borders corresponding to the 25^th^ and 75^th^ percentiles of the data. Different letters denote significance at *P* < 0.05 (one-way ANOVA followed by Tukey’s post hoc test). (C) Hydrogen peroxide accumulation in Arabidopsis plants in response to WLS. Representative data of 6 independent replicates is shown as box and whisker plot with borders corresponding to the 25^th^ and 75^th^ percentiles of the data. Different letters denote significance at *P* < 0.05 (one-way ANOVA followed by a Tukey’s post hoc test). Abbreviations: CT, control; H_2_DCFDA, 2’,7’-dichlorodihydrofluorescein diacetate; H_2_O_2_, hydrogen peroxide; ROS, reactive oxygen species; WLS, waterlogging stress.

### Waterlogging stress triggers a systemic hydraulic wave response and causes the transient opening of stomata

Plants respond to different treatments that abruptly alter the water pressure in their vascular system with a hydraulic wave (*e.g.,* wounding; Kloth and Dicke, 2022; Grenzi et al., 2023; Gao et al., 2023). As the sudden application of WLS could potentially activate a hydraulic wave in plants due to an increase in the water pressure around the root system, we measured the systemic hydraulic wave in plants subjected to WLS (Zimmermann et al., 2013; Fichman and Mittler, 2021). As shown in Figure 3A, WLS resulted in the triggering of a rapid systemic hydraulic wave response that was detected in shoots within 5 minutes of WLS application to the root system. Interestingly, the application of a sudden WLS to plants also resulted in a transient stomatal opening response that started at about 1 min following the application of WLS and lasted for about 10 min (Figure 3B). To examine whether the hydraulic wave (Figure 3A), and the transient stomatal opening response (Figure 3B), triggered by WLS were linked, we pre-treated plants with ABA that caused stomata to close and applied WLS. As shown in Figures 3A and 3B, pretreatment of plants with ABA (50 μM) 150 min before WLS suppressed the hydraulic wave response as well as the transient stomatal response of plants to the sudden WLS treatment.

**Figure 3.**
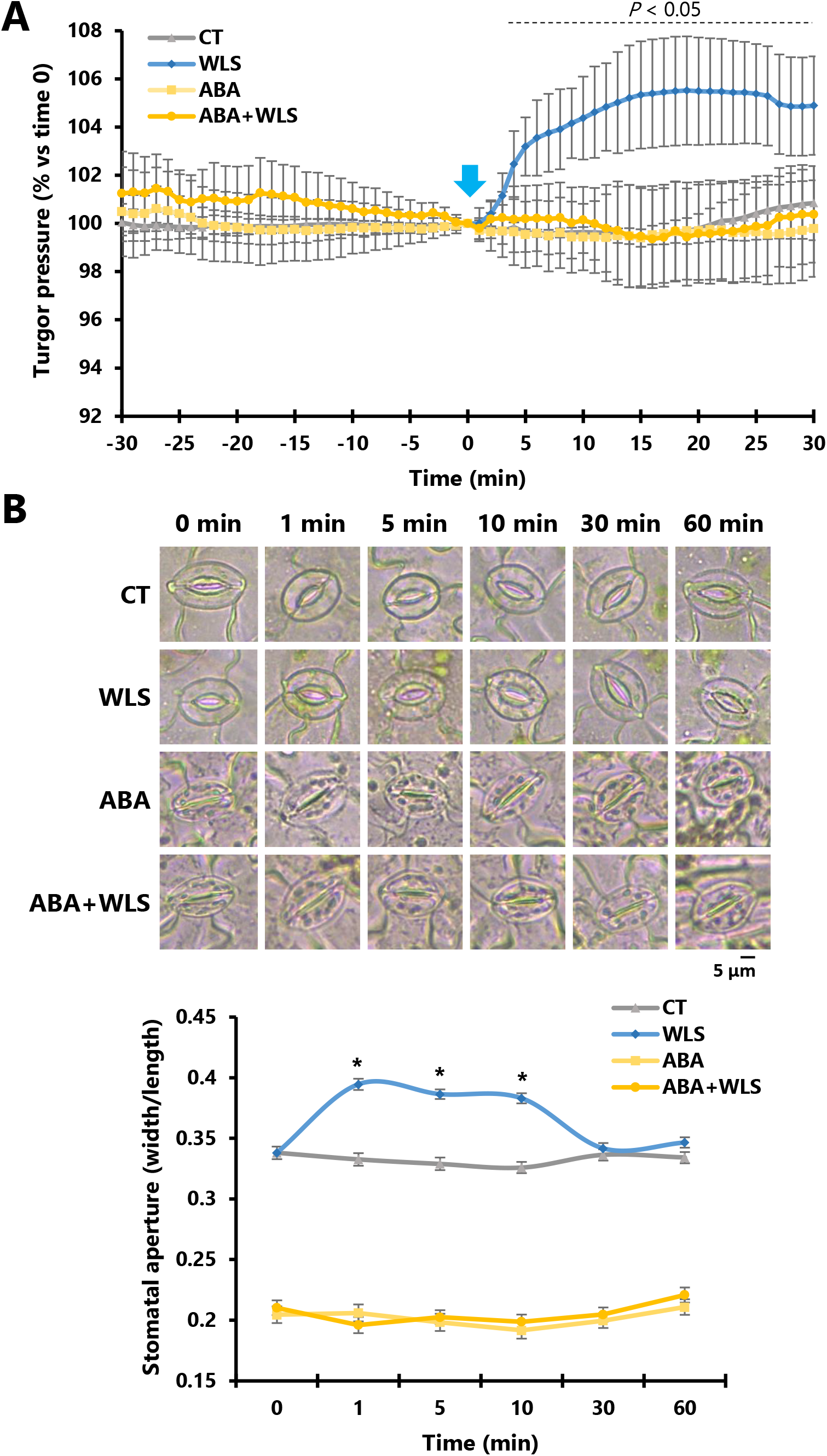
Systemic changes in hydraulic pressure and stomatal responses in wild type plants subjected to waterlogging stress (WLS). (A) Continuous systemic leaf turgor pressure measurements of WT plants from 30 min prior (−30 min) to 30 min post (30 min) WLS (applied at 0-min time; indicated with an arrow). Graph shows hydraulic pressure in control (CT) plants and plants subjected to WLS. Water (CT) or ABA (50 µM; ABA and ABA+WLS) were applied to plants by spraying 150 min before WLS application. Hydraulic pressure is represented as the percentage of the initial measured turgor pressure at 0 min. Statistical analysis was performed with a two-sided Student’s t test (\**P* < 0.05; n = 12). All experiments were repeated at least 5 times with 3 plants per treatment. (B) Systemic stomatal aperture response of Arabidopsis to WLS. Representative images of stomata from control (CT) and plants treated (50 µM; ABA and ABA+WLS) or untreated with ABA (WLS; sprayed with water) 150 min before WLS are shown on left, and line graphs showing stomatal aperture measurements at 1-, 5-, 10-, 30- and 60-min following WLS application are shown on right. Results were obtained using at least 20 different plants for each time and treatment (means ± SE, n = 500; \**P* < 0.05). Stomatal aperture data was compared with control plants at each time point using two-sided Student’s t test (\**P* < 0.05). Scale bar in (B) represents 5 µm. Abbreviations: ABA, abscisic acid; CT, control; WLS, waterlogging stress.

### The WLS-triggered ROS wave is dependent on RBOHD, GLR3.3GLR3.6 and PIP2;1 function

The systemic ROS wave response of Arabidopsis to a local treatment of excess light stress or wounding was previously shown to depend on the function of different proteins such as RBOHD, GLR3.3GLR3.6, PLASMODESMATA LOCALIZED PROTEIN 5 (PDLP5) and/or PIP2;1 (Fichman et al., 2021; Fichman and Mittler, 2021). To test whether the WLS-triggered ROS wave (Figure 2A) is also dependent on RBOHD function, we applied a drop of the broad-range oxidase and RBOH inhibitor diphenyleneiodonium (DPI, 50 µM) or water, in agarose, to the middle point between the root system and the shoot (just above the peat soil level, as described in Devireddy et al., 2018), 30 min prior to subjecting plants to WLS. In addition, we compared the response to WLS between wild type and the *rbohD* Arabidopsis mutant. As shown in Figure 4, pretreatment of plants with a drop of agar containing DPI prior to WLS (Figure 4A), or treatment of the *rbohD* mutant (Figure 4B) with WLS resulted in the suppression of the ROS wave response induced by WLS in Arabidopsis. A similar result was found when whole plants were fumigated with DPI 30 min prior to the application of WLS (Supplementary Figure S1). These findings suggest that RBHOD function is required for the WLS-induced ROS wave response.

**Figure 4.**
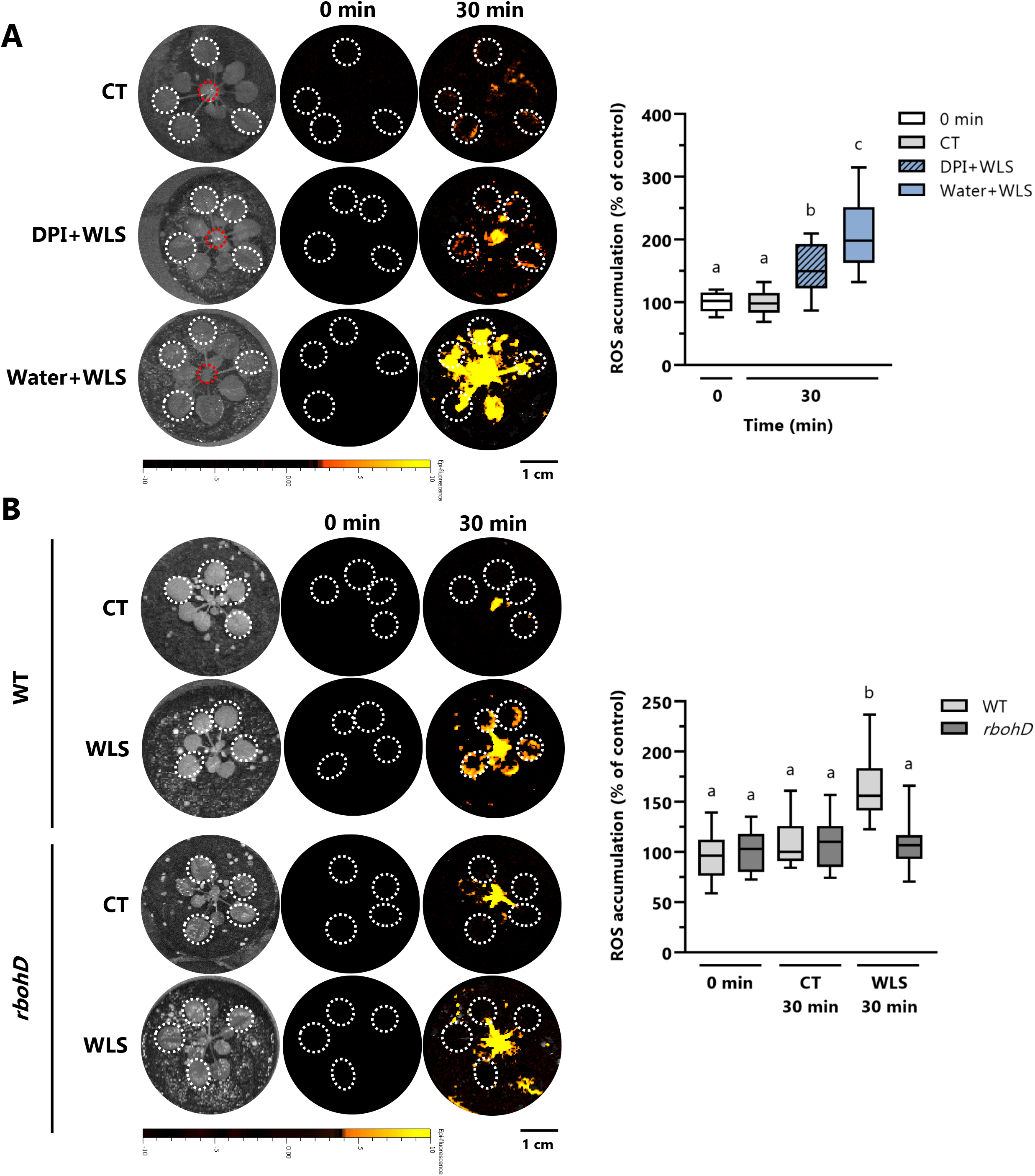
Suppression of the systemic ROS signal triggered by waterlogging stress (WLS). (A) The NADPH oxidase inhibitor DPI (50 µM) or water were applied in a drop of agarose to the middle point between the root system and the shoot and plants (indicated with a dashed red circle) and plants were fumigated with H_2_DCFDA for 30 min before applying WLS. Representative time-lapse images of whole plant ROS accumulation in treated and untreated *Arabidopsis thaliana* plants are shown alongside bar graphs of combined data from all plants used for the analysis at the 0- and 30-min time points. Data is shown as box and whisker plots with borders corresponding to the 25^th^ and 75^th^ percentiles of the data. (B) Time-lapse imaging (left) and bar graphs (right) of combined ROS accumulation data in WT and the *rbohD* mutant in response to a sudden WLS treatment. Regions of interest are indicated by white ovals in the images in (A) and (B). Magnification bars in (A) and (B) are 1 cm. Different letters denote significance at *P* < 0.05 (ANOVA followed by a Tukey’s post hoc test). All experiments were repeated at least three times with eight plants per repeat. Abbreviations: CT, control; WLS, waterlogging stress; DPI, diphenyleneiodonium; H_2_DCFDA, 2’,7’-dichlorodihydrofluorescein diacetate; RBOHD, NADPH/respiratory burst oxidase protein D; WT, wild type.

To determine the role of GLR3.3GLR3.6, PDLP5, and PIP2;1, previously found to be involved in regulating the ROS wave response to injury or excess light stress (Fichman et al., 2021; Fichman and Mittler, 2021), in mediating the WLS-triggered ROS wave response, we subjected wild type and *glr3.3glr3.6*, *pdlp5*, *pip2;1*, and *pip1;4* mutants to WLS and measured their systemic ROS wave response. As shown in Figure 5A, the function of GLR3.3GLR3.6 and PIP2;1 was required for the WLS-induced ROS wave response, while the function of PIP1;4 and PDLP5 was not.

**Figure 5.**
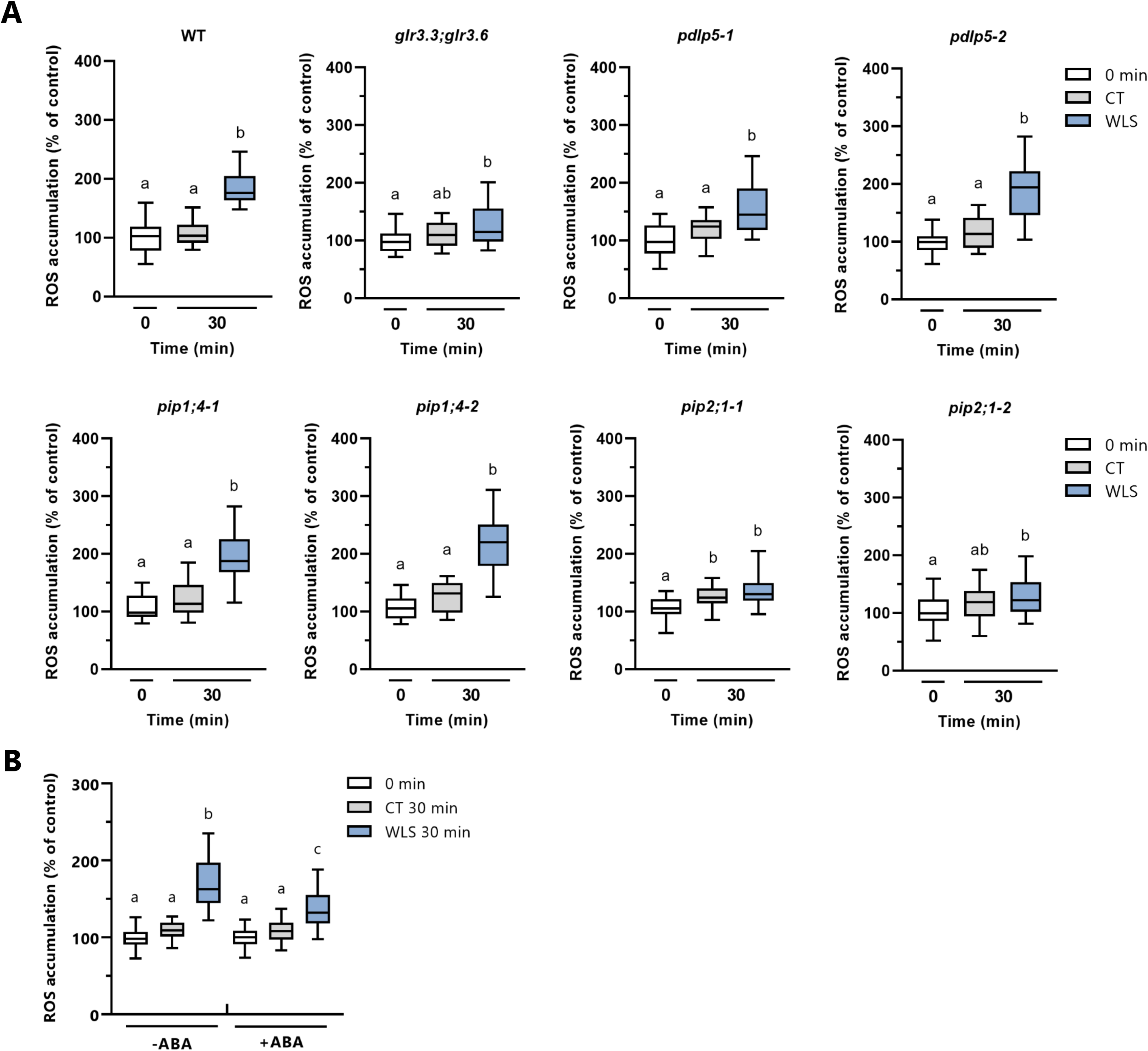
GLR3.3;GLR3.6 and PIP2;1 are required for waterlogging stress (WLS)-induced rapid systemic ROS signaling in Arabidopsis. (A) ROS accumulation was measured in wild type (WT) and the *glr3.3;glr3.6, pdlp5-1*, *pdlp5-2*, *pip1;4-1*, *pip1;4-2*, *pip2;1-2* and *pip1;1-2* mutants in response to a sudden WLS treatment. (B) The effect of pretreatment with ABA on the systemic ROS wave response of wild type plants to WLS. Bar graphs of combined data from all plants used for the analysis at the 0- and 30-min time points are shown. All experiments were repeated at least three times with eight plants per repeat. Representative data is shown as box and whisker plots with borders corresponding to the 25^th^ and 75^th^ percentiles of the data. Different letters denote significance at *P* < 0.05 (ANOVA followed by a Tukey’s post hoc test). Abbreviations: ABA, abscisic acid; CT, control; glr, glutamate receptor-like; pdlp5, plasmodesmata localized protein 5; pip1, plasma membrane intrinsic protein 1; pip2, plasma membrane intrinsic protein 2; WLS, waterlogging stress; WT, wild type.

As, pre-treatment of plants with ABA blocked the transient stomatal opening response of plants, as well as the hydraulic wave response, to WLS (Figure 3), we also tested whether pre-treatment with ABA will block the ROS wave in response to WLS. As shown in Figure 5B, pre-treatment of plants with ABA suppressed the ROS wave response to WLS.

### Rapid systemic transcriptomic responses to WLS in wild type and the *rbohD* mutant

The transcriptomic response of Arabidopsis to anoxia, hypoxia, submergence, or simulated WLS was previously studied in Arabidopsis (Liu et al., 2005; Hsu et al., 2011; Lee et al., 2011; Licausi et al., 2011; Pucciariello et al., 2012; Hsu et al., 2013; Tsai et al., 2014; van Veen et al., 2016; Giuntoli et al., 2017; Liu et al., 2017; Bui et al., 2020; Yang et al., 2022). However, most of these studies did not focus on rapid transcriptomic responses and/or did not use soil or peat for plant growth. To examine the systemic response of shoots from peat soil-grown plants subjected to WLS, we conducted a transcriptomic analysis of wild type plants subjected to a 0-, 10-, 30-, and 60-min of WLS (Figure 6; Table 1; Supplementary Tables S1-S9). In addition, we tested the expression of several known hypoxia- and ROS-related transcripts by qPCR at 0- and 60-min post WLS initiation, to ascertain that the waterlogging treatment we applied to plants induced systemic responses to hypoxic conditions (Supplementary Figure S2). Waterlogging stress caused the altered expression of over 2,400, 3,500, and 6,300 transcripts within 10-, 30-, and 60-min of stress initiation respectively, with over 300, 800, and 3,800 transcripts uniquely altered in each of these time points (Figure 6A; Supplementary Tables S1-S6). Collectively, transcripts altered in systemic tissues in response to WLS contained a high representation of stress-, stimuli-, and anoxia-response transcripts, as well as transcripts involved in hormone, cell communication, and biotic and abiotic responses (Figure 6B; Supplementary Table S7). A significant overlap was found between the transcripts identified by our study in systemic tissues of plants subjected to WLS (Figure 6A) and transcripts identified by several other studies (Bui et al., 2020; Tamura and Bono, 2022) in whole plants subjected to hypoxia or submergence (Figure 6C; Supplementary Table S8). Less overlap was nevertheless found with transcriptomic data obtained from shoots of plants subjected to anoxic conditions or a simulated WLS (Tsai et al., 2014; van Veen et al., 2016); potentially due to the different conditions and time points used in the two studies (Figure 6D; Supplementary Table S8).

**Figure 6.**
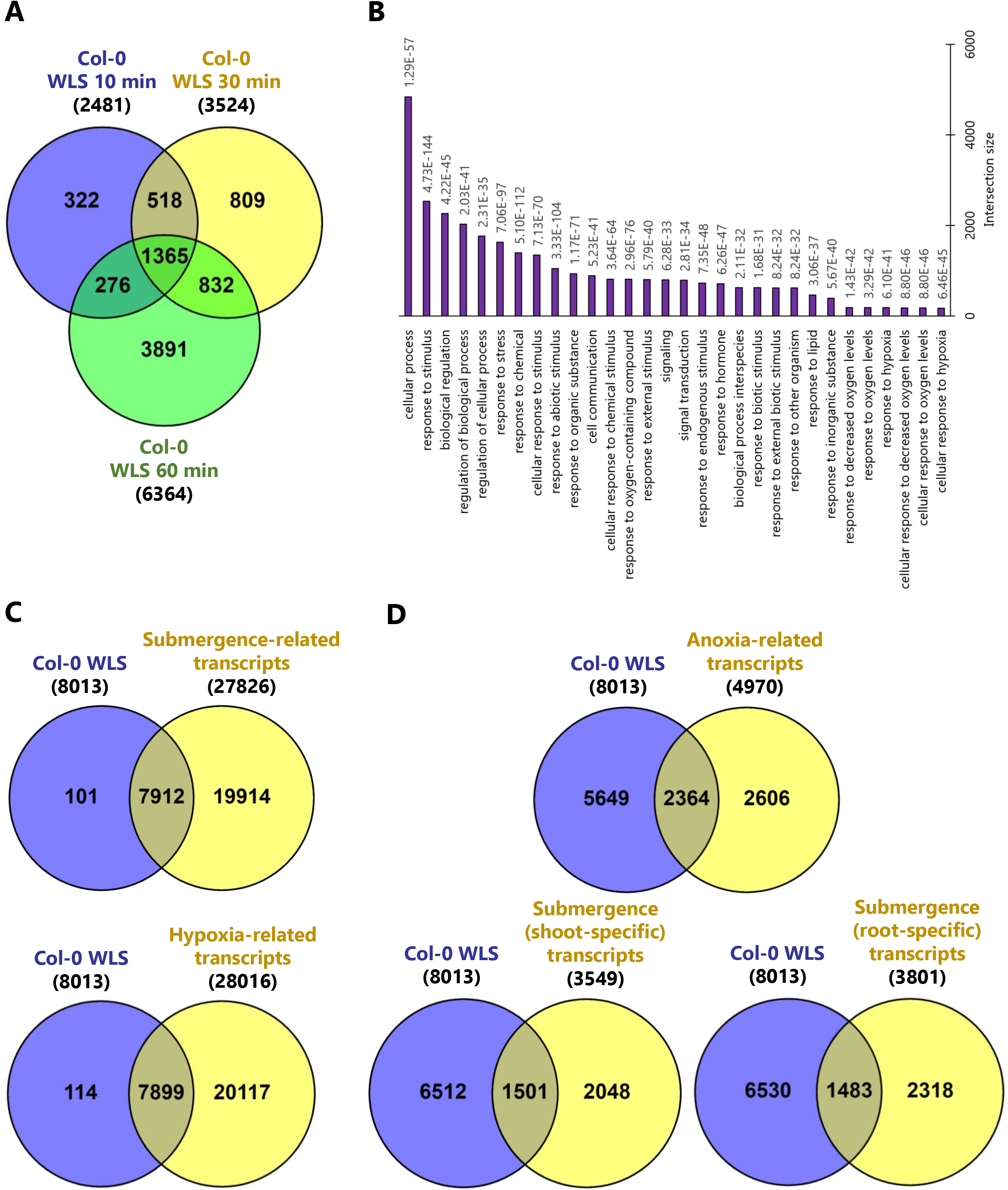
Transcriptomic analysis of the systemic response of wild type and *rbohD* plants subjected to waterlogging stress (WLS). (A) Venn diagram showing the overlap in transcripts significantly altered in wild type (WT) after 10-, 30- and 60-min of WLS compared to 0 min. The 30 most statistically significant categories found in Biological Process (BP) from Gene ontology (GO) annotation of all the transcripts (8,013) altered in systemic tissues in response to WLS in WT. (C) and (D) Comparison between the transcripts altered in WT in response to WLS in this study (8,013) and transcriptomic data from previous studies. References used for the comparisons in (C) are Bui et al., 2020 (submergence-related transcripts) and Tamura and Bono 2022 (hypoxia-related transcripts). In (D) the references used are Tsai et al., 2014 (anoxia-related transcripts) and van Veen et al., 2016 (shoot and root-specific transcripts). Abbreviations: WLS, waterlogging stress.

**Table 1.** Representation of stress-, hormone-, and reactive oxygen species (ROS)-response transcripts in the different groups of transcripts significantly altered in WT and rbohD (RBOHD-dependent transcripts) plants in response to waterlogging stress (WLS). The transcripts significantly altered in plants in response to stress, hormone, and ROS (left column) from categories ‘cold’, ‘drought’, ‘heat’, ‘salt’, ‘wounding’, ‘ozone’, ‘pathogen’, ‘photosynthesis’, ‘TFs’, ‘ROS-responsive’, ‘O_2_^-^’, ‘^1^O_2_’, ‘H_2_O_2_’, ‘ROS wave’, ‘ABA’, ‘BL’, ‘CK’, ‘GA’, ‘IAA’, ‘JA’, ‘SA’ were obtained from Zandalinas et al., (2019, 2021a); the transcripts included in ‘hypoxia core’ were taken from Mustroph et al., 2009; and the transcripts in ‘hypoxia’ and ‘ET’ categories were obtained from the GO database using as filter the term IDs GO:0071456, GO:0036293, GO:0036294, GO:0001666 for ‘hypoxia, and GO:0009723, GO:0009873, GO:0071369. The percent of transcripts included in each of these sets (stress-, hormone-, and ROS-response transcripts), found in the datasets obtained in the current study (top row) is shown. The total number of transcripts included in each of the stress-, hormone-, and ROS-response transcript datasets is shown in brackets (left column). Values above 50% are highlighted in bold. Abbreviations: ABA, abscisic acid; ET, ethylene; BL, brassinolide; CK, cytokinins; GA, gibberellic acid; IAA, indole-3-acetic acid; JA, jasmonic acid; RBOHD, NADPH/respiratory burst oxidase protein D; RBOHD-dep, RBOHD-dependent transcripts; SA, salicylic acid; TFs, transcription factors; WLS, waterlogging stress; WT, wild type.

|  | WT WLS<br>10 min<br>(2,481) | WT WLS<br>30 min<br>(3,524) | WT WLS<br>60 min<br>(6,364) | WT WLS<br>10+30+60<br>min<br>(8,013) | RBOHD-dep<br>(1,806+<br>3,028) |
| --- | --- | --- | --- | --- | --- |
| **Cold** (622) | 49.0 | **59.5** | **63.7** | **76.4** | 24.0 |
| **Hypoxia** (358) | 40.8 | 47.2 | **51.1** | **61.7** | 20.1 |
| **Drought** (3,382) | 16.9 | 25.3 | 39.1 | 49.6 | 26.5 |
| **Heat** (1,716) | 19.6 | 25.3 | 36.2 | 46.4 | 24.7 |
| **Salt** (1,614) | 26.1 | 34.9 | 35.5 | 48.6 | 25.9 |
| **Wounding** (3,882) | 22.6 | 32.8 | 44.3 | **56.0** | 24.6 |
| **Ozone** (764) | 29.7 | 39.8 | 47.0 | **58.9** | 27.4 |
| **Pathogen** (581) | 23.1 | 31.2 | 35.3 | 47.0 | 27.9 |
| **Photosynthesis** (67) | 25.4 | 34.3 | 23.9 | **52.2** | 37.3 |
| **TFs** (524) | 6.3 | 7.1 | 8.4 | 11.3 | 6.5 |
| **ROS-responsive** (1,282) | 37.8 | 48.8 | **52.7** | **67.4** | 25.0 |
| **O<sub>2</sub><sup>-</sup>** (286) | 37.4 | 49.0 | 46.2 | **62.9** | 29.4 |
| **<sup>1</sup>O<sub>2</sub>** (297) | 33.0 | 47.8 | 44.1 | **62.6** | 25.3 |
| **H<sub>2</sub>O<sub>2</sub>** (956) | 43.2 | **52.6** | **58.5** | **72.1** | 23.7 |
| **ROS wave** (82) | **63.4** | **74.4** | **67.1** | **84.1** | 26.8 |
| **Hypoxia core** (49) | 36.7 | 40.8 | 40.8 | 49.0 | 14.3 |
| **ET** (255) | 13.3 | 17.3 | 38.0 | 43.1 | 27.5 |
| **ABA** (1,460) | 19.7 | 28.9 | 45.9 | **56.6** | 27.3 |
| **BL** (276) | 37.7 | 49.3 | **56.5** | **70.3** | 25.7 |
| **CK** (335) | 20.0 | 23 | 44.8 | **52.8** | 24.8 |
| **GA** (43) | 41.9 | 46.5 | 39.5 | **60.5** | 30.2 |
| **IAA** (436) | 30.7 | 39.9 | 48.4 | **59.4** | 31.2 |
| **JA** (3,877) | 21.2 | 30.8 | 44.4 | **55.8** | 26.4 |
| **SA** (217) | 21.7 | 34.6 | **61.3** | **68.2** | 22.1 |

When comparing the transcripts significantly altered in our dataset (Figure 6A) with different sets of transcripts significantly altered in plants subjected to different stresses, hormone treatments, or ROS (Zandalinas et al., 2019, 2021a), it was found that many cold-, hypoxia-, wounding-, and ozone-response transcripts are altered in their expression in shoots of plants subjected to WLS (Table 1; Supplementary Table S9). In addition, many ROS- and/or ROS wave-response transcripts, previously identified by other studies (Zandalinas et al., 2019) were altered in their expression in shoots of plants subjected to WLS.

To examine what proportion of the systemic transcriptomic response of Arabidopsis shoots to WLS is dependent on RBOHD, we conducted an RNA-Seq analysis on wild type and *rbohD* mutant plants subjected to WLS for 0- and 60-min (Supplementary Tables S10-S14). As shown in Figure 7A, over 1,800 transcripts that were altered in wild type plants in response to WLS were not altered in the *rbohD* mutant. In addition, over 3,000 transcripts were specifically altered in the *rbohD* mutant, but not wild type (Figure 7A; Supplementary Table S11). As shown in Table 1 (Supplementary Table S12), over 20% of all stress-, ROS-, hypoxia-, and hormone-response transcripts were altered in their expression in the combined group of RBOHD-dependent transcripts (1,806+3,028). RBOHD-dependent transcripts were also enriched in abiotic-, biotic-, ethylene-, and ABA-response transcripts, and transcripts involved in organic and metabolic processes (Figure 7B; Supplementary Table S13). In addition, as shown in Figure 7C, 27, 29, and 28% of all transcripts altered in their expression at 10-, 30- and 60-min, respectively, were RBOHD-dependent (Supplementary Table S14).

**Figure 7.**
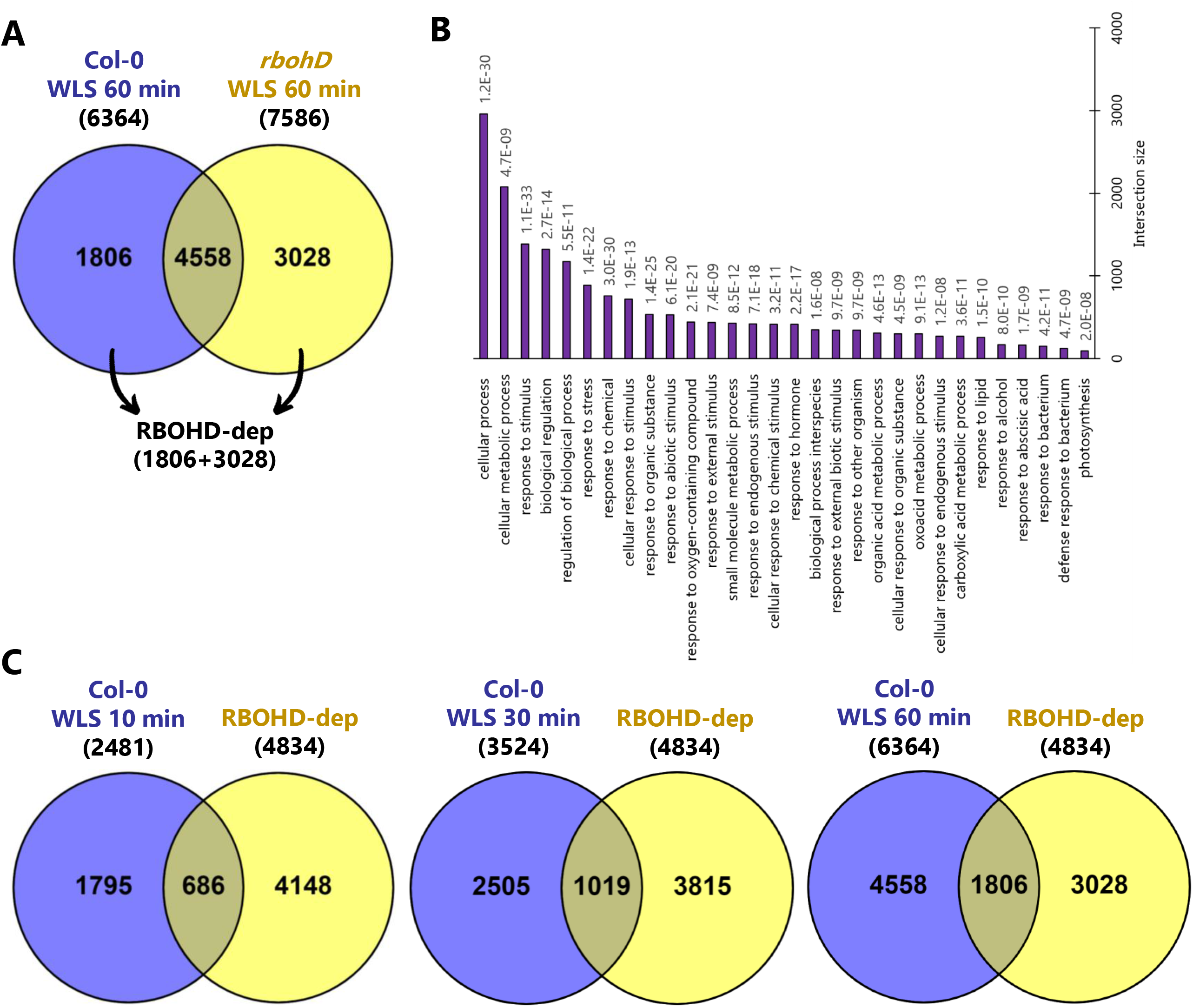
RBOHD-dependent systemic transcriptomic responses to WLS. (A) Overlap between transcripts significantly altered in WT and *rbohD* plants following 60 min of WLS treatment. (B) The 30 most statistically significant categories found in Biological Process (BP) of the GO annotation for the RBOHD-dependent transcripts (1,806+3,028) at time 60 min. (C) Venn diagrams showing the overlap between the RBOHD-dependent transcripts at 60 min (1,806+3,028) and the transcripts altered in WT after 10-, 30- and 60-min of WLS. Abbreviations: RBOHD, NADPH/respiratory burst oxidase protein D; RBOHD-dep, RBOHD-dependent transcripts; WLS, waterlogging stress.

## DISCUSSION

Plants can respond to stress within seconds to minutes of stress initiation triggering multiple molecular, metabolic, and physiological mechanisms (Suzuki et al., 2015; Choudhury et al., 2018; Devireddy et al., 2018; Kollist et al., 2019). Rapid responses to stress are especially important for plants experiencing the initiation of WLS in the field or in nature; that could potentially be followed by partial or complete submergence if the advance of flood water or downpour will not stop (Voesenek and Bailey-Serres, 2015; Loreti et al., 2016; Pucciariello and Perata, 2017; Sasidharan et al., 2018, 2021). Here we report that a sudden WLS treatment of Arabidopsis is followed by rapid changes in the expression of many hypoxia-response transcripts in shoots in a time-dependent manner (Figure 6; Table 1). This finding reveals that plants rapidly activate hypoxia acclimation pathways in shoots (while still under aerobic conditions) potentially in anticipation of an impending partial or complete submergence (that would be accompanied by hypoxia stress). This response is analogous to other rapid systemic responses, *e.g.,* to pathogens, insects, and/or abiotic stresses such as excess light or temperature extremes (Kollist et al., 2019), and represents an important evolutionary advantage for multicellular organisms that are sessile.

Previous studies conducted with trees and other plants, including crop plants, identified stomatal responses, changes in hydraulic pressure (in the roots and stems), and changes in transcriptomic responses, as playing an important role in plant responses and acclimation to WLS (*e.g.,* Sellin, 2001; Rodríguez-Gamir et al., 2011; Liu et al., 2012; Striker et al., 2014; Jurczyk et al., 2016; Gasch et al., 2016; Martínez-Arias et al., 2020; Repo et al., 2021; Liu et al., 2022). These studies were however conducted in plants subjected to longer WLS periods and did not address rapid responses that occur within minutes of WLS initiation. In our work we demonstrate that stomatal, hydraulic, and transcriptomic responses occur in systemic tissues of Arabidopsis plants within 1-10 min of WLS application (Figures 2, 3, and 6), revealing that many important responses to WLS might occur much earlier than previously thought. The overlap between many of the responses identified by our study and previous studies (stomatal, hydraulic, and transcriptomics), suggests that the early changes we identified in our study play an important role in plant acclimation to WLS.

Cellular responses to hypoxia and/or anoxia have been extensively studied, and RBOHD was found to play a key role in ROS accumulation during the initial stages of anoxia stress (as well as in plant acclimation to anoxia; Liu et al., 2017). Our study reveals that in addition to controlling local responses to anoxia (Liu et al., 2017), RBOHD could also be involved in controlling systemic responses to WLS (Figures 4, 7; Table 1). The activation of RBOHD in roots following the sudden WLS treatment could therefore propagate in a cell-to-cell fashion from the roots all the way to the shoots and trigger the expression of many different anoxia and hypoxia acclimation mechanisms in the shoots (Figures 2, 4, 6; Table 1). Our findings that the application of a drop of DPI to the middle point between the roots and the shoots suppresses the WLS-induced ROS wave in shoots (Figure 4A), strongly support this possibility. As with previous studies of rapid systemic ROS signaling in Arabidopsis in response to different abiotic stresses, the WLS-triggered ROS wave required the function of GLR3.3GLR3.6 and PIP2;1 (Figure 5A; Fichman et al., 2021; Fichman and Mittler, 2021). In contrast, PDLP5, that was previously found to be required for systemic responses to wounding or excess light stress (Fichman et al., 2021; Fichman and Mittler, 2021), appeared to not be required for this response (Figure 5A). These findings suggest that the ROS wave induced by WLS (Figure 2A) is mediated via a somewhat different molecular mechanism than that triggered in response to excess light or wounding in Arabidopsis (Figure 5A; Fichman et al., 2021; Fichman and Mittler, 2021). Further studies are required to address the molecular mechanisms regulating the ROS wave response to WLS.

We previously reported that the calcium, ROS, and hydraulic waves are integrated in Arabidopsis during systemic responses to wounding (applied to a single leaf) and require the function of GLR3.3GLR3.6 (Fichman et al., 2021; Fichman and Mittler, 2021). Although under WLS the ROS wave was dependent on GLR3.3GLR3.6 function (Figure 5A), it is not clear whether the calcium and hydraulic waves are also dependent on these calcium-permeable channels. In addition, the function of CAX1, previously found to control calcium and ROS signaling under submergence and anoxia stresses (Yang et al., 2022), in mediating the ROS wave during WLS is unclear. One possibility is that in roots, under waterlogging-mediated anoxia stress, CAX1 is required for ROS production by RBOHD (Yang et al., 2022), while in shoots, under aerobic conditions, GLR3.3GLR3.6 are required for ROS production by RBOHD (Figure 5A; Fichman et al., 2021; Fichman and Mittler, 2021). As calcium signaling plays such a key role in triggering abiotic and biotic responses in plants (*e.g.,* Luan and Wang, 2021), further studies are needed to address the potential contribution of these calcium signaling mechanisms, as well as others, to the overall systemic response of plants to WLS.

Hydraulic waves are thought to play an important role in the systemic response of plants to different stresses (Kloth and Dicke, 2022; Grenzi et al., 2023). The changes in water pressure propagating through the vascular system of plants were proposed to trigger mechanosensory proteins that translate the hydraulic signal into a calcium signal and subsequently a ROS accumulation response in systemic tissues (Gilroy et al., 2016; Kloth and Dicke, 2022). As the systemic response of Arabidopsis to WLS involves a rapid systemic hydraulic response (Figure 3A), that was followed by systemic calcium and ROS responses (Figure 2), it is possible that these three waves are interlinked in Arabidopsis during waterlogging stress. This possibility is also supported by the finding that pre-treatment of plants with ABA, that suppressed the hydraulic wave (Figure 3A), also suppressed the ROS wave (Figure 5B). The association between the hydraulic, calcium, and ROS waves should be addressed in future studies using experimental systems like the one presented in this study, using for example mechanosensory mutants such as the MECHANOSENSITIVE ION CHANNEL LIKE 2 or 3 (*msl2* or *msl3*) mutants, previously found to be required for the propagation of the ROS wave (Fichman et al., 2021, 2022). The association between the stomatal opening response, ABA, and the hydraulic wave (Figure 3) should also be pursued in future studies as it suggests that the degree of stomatal aperture openness at the time WLS occurs might impact hydraulic waves and other systemic responses to this stress. In this respect, it should be mentioned that hydraulic waves were recently proposed to control glutamic acid release from cells during systemic responses to wounding, controlling the systemic calcium wave (Grenzi et al., 2023), as well as linked to the mobilization of glucohydrolases, which are important for the regulation of electrical signals during wound responses (Gao et al., 2023). Hydraulic waves might therefore play a central role in regulating systemic responses in plants, and elucidating their function requires further studies.

The potential dependence of systemic hydraulic responses to WLS on stomatal aperture (Figure 3) could play an important role during conditions of stress combination (Zandalinas and Mittler, 2022). It was previously shown that in combination with other stresses, such as heat stress (that is becoming a major problem worldwide due to global warming; Bailey-Serres et al., 2019; Zandalinas et al., 2021b; Masson-Delmotte et al., 2021), WLS can become significantly more lethal to some crops (*e.g.,* Lin et al., 2015a; Lin et al., 2015b; Zhen et al., 2020; Shao et al., 2022), highlighting the importance of studying plant responses to complex conditions of stress combination (Zandalinas et al., 2021b; Zandalinas and Mittler, 2022). As rapid systemic responses to different stresses could be conflicting during stress combination (Zandalinas et al., 2020a), more studies are needed to address the effects of heat and other stresses on the rapid systemic responses of plants to WLS. This is especially important as WLS causes rapid alterations in stomatal aperture (Figure 3B), and stomatal responses to different stresses, occurring during stress combination, were shown to be contradicting (Zandalinas and Mittler, 2022).

Although the 10 min WLS treatment described in the current study resulted in a decrease in oxygen levels around the plant root system (Figure 1B), and the systemic transcriptomic response at 10 min included hypoxia-related transcripts (Figure 6; Table 1), the rapid systemic transcriptomic response of Arabidopsis to the sudden WLS treatment observed in our study might in fact reflect a combination of different systemic signals; some to hypoxia conditions (Figure 1B), some to changes in hydraulic pressure (Figure 3A), and some to increase in ethylene levels around the root system (Hsu et al., 2011; Tsai et al., 2014). One way to dissect the different causes of the observed rapid systemic transcriptomic response could be to study it in mutants deficient in hypoxia responses (*e.g.,* mutants impaired in GROUP VII ETHYLENE RESPONSE FACTORs; Gasch et al., 2016), mutants deficient in ethylene signaling (Myers et al., 2023), and mutants deficient in hydraulic wave responses (Fichman and Mittler, 2021). Such studies could distinguish between hypoxia-, ethylene-, or hydraulic-driven transcriptomic responses. In this respect it is important to note that in contrast to the large number of hypoxia-response transcripts accumulating at 10 min following WLS initiation, the number of ethylene-response transcripts accumulating at 10 min was not high (Table 1). This finding, together with the oxygen level measurements around the root system at 10 min (Figure 1B), suggest that the systemic hypoxia-related response observed in our study at 10 min is indeed at least partially linked to the induction of hypoxic conditions at the root system following WLS.

## MATERIALS AND METHODS

### Plant material and growth conditions

Seeds of *Arabidopsis thaliana* Col-0 (cv. Columbia-0) and the mutants *rbohD* (Torres et al., 2002), *glr3.3glr3.6* (Mousavi et al., 2013; Nguyen et al., 2018), *pdlp5* (two independent alleles, Fichman et al., 2021), *pip2;1* (two independent alleles, Rodrigues et al., 2017; Fichman et al., 2021) and *pip1;4* (two independent alleles, Fichman et al., 2021) were germinated and grown on peat pellets (Jiffy-7; Jiffy International, Kristiansand, Norway), under controlled conditions of 10-hour/14-hour light/dark regime, 50 µmol photons s^−1^ m^−2^, and 21°C for 4 weeks.

### Waterlogging stress application

To induce WLS, we grew plants as described above for 4 weeks, allowed the water content of the peat pellets to reach 54 ± 2% of complete peat soil water saturation (each peat pellet was first weighted when it was fully saturated with water and then weighted regularly until 54 ± 2 % of fully saturated weight was achieved). Plants were then treated or untreated with the different dyes for imaging as described below, placed in a tray that allowed watering to saturation inside the imager, or back under the growth light, as described below, rapidly watered until the water levels reached all the way to the top of the peat soil (97 ± 5% water capacity measured 60 min following watering), and subjected to imaging or other analyses as described below (Figure 1A). Oxygen levels around the root system were measured before and during the WLS with the Bante821 portable dissolved oxygen meter (Bante Instruments, Shanghai, China).

### Whole-plant imaging of ROS and calcium levels

As previously described (Fichman et al., 2019; Fichman et al., 2020b; Zandalinas et al., 2020a,b; Fichman and Mittler, 2021), plants were fumigated for 30 min with 50 µM 2′,7′-dichlorofluorescein diacetate (H_2_DCFDA; Millipore-Sigma, St. Louis, MO, USA) for ROS imaging, or with 4.5 µM Fluo-4-AM (Becton, Dickinson and Company, Franklin Lakes, NJ, USA) for calcium imaging, using a nebulizer (Punasi Direct, Hong Kong, China) in a glass container. To inhibit ROS propagation, diphenyleneiodonium (DPI, 50 µM, Millipore-Sigma, St. Louis, MO, USA) or water were applied in a drop of 0.3% agarose to the midpoint of the rosette that connects the shoot to the root system 30 min before WLS, as described by Fichman et al. (2019). When plants were also fumigated with a solution containing DPI at a final concentration of 50 µM together with the dye H_2_DCFDA as described above. Following fumigation, plants were subjected to a sudden WLS as described above, a black plastic mask with holes was placed above the tray containing the water to avoid the background of water autofluorescence, and imaging using the IVIS Lumina S5 fluorescence imager (PerkinElmer, Waltham, MA, USA). Fluorescence images (excitation/emission 480 nm/520 nm) were acquired every minute for 60 min as described by Fichman et al. (2019). Images were analyzed with Living Image 4.7.2 software (PerkinElmer) as explained previously (Fichman et al., 2019). All experiments were repeated at least three times each with 8 plants.

### H_2_O_2_ detection

Hydrogen peroxide quantification in systemic leaves was performed using Amplex-Red (10-acetyl-3,7-dihydroxyphenoxazine; ADHP; Thermo Fisher Scientific, Waltham, MA, USA) as described by Fichman et al. (2022). Systemic leaves from control and WLS-treated plants were immediately frozen and ground to fine powder, resuspended in 50 µL 0.1 M trichloroacetic acid (TCA; Thermo Fisher Scientific, Waltham, MA, USA), and centrifuged for 15 min at 12,000g, 4°C. The supernatant was buffered with 1 M phosphate buffer pH 7.4, and the pellet was dried and used for dry weight calculation. H_2_O_2_ quantification in the supernatant was performed according to the MyQubit-Amplex-Red Peroxide Assay manual (Thermo Fisher Scientific, Waltham, MA, USA), using a calibration curve of H_2_O_2_ (Thermo Fisher Scientific, Waltham, MA, USA) as described by Fichman et al. (2022).

### Hydraulic pressure measurements

Changes in systemic leaf turgor pressure following WLS were recorded using the ZIM-probe system (Yara International ASA, Oslo, Norway; Zimmermann et al., 2013), as described by Fichman and Mittler (2021). Briefly, a single leaf of four-week-old plants was connected to two magnetic probes that included a pressure sensor between them (Zimmermann et al., 2013). The turgor pressure force against the magnetic pressure was recorded and transmitted to a receiver every minute. Abscisic acid (ABA) treatment was performed by spraying ABA (50 µM; Millipore-Sigma, St. Louis, MO, USA) on the entire rosette 30 min prior to connecting leaves to the magnetic probes. Control plants were simultaneously sprayed with distilled water. Following magnetic probe attachment, the system was allowed to stabilize for 2 h and plants were subjected to the WLS treatment. Pressure values were recorded for an additional 60 min following the stress application. Hydraulic pressure was calculated as the percentage of the initial measured turgor pressure at 0 min, which is the pressure in the leaf right before the stress application. All experiments were performed between 9 AM and 1 PM. Each dataset includes average and SE of 6 to 12 biological repeats.

### Measurements of stomatal aperture

A thin layer of nail polish (450B clear nail protector-Wet N Wild; Markwins Beauty Products, CA, USA) was applied to the abaxial surface of the leaf avoiding the major veins. The nail polish was left to dry for approximately ten minutes and then peeled off with tweezers. Impressions were mounted pointing upward with double-sided tape (Scotch) on a microscope slide. Stomata images were captured with an EVOS XL microscope (Invitrogen by Thermo Fisher Scientific, Waltham, MA, USA). Both width and length of stomatal aperture were measured using IMAGE J (https://imagej.nih.gov/ij). Stomatal aperture was calculated as a ratio of stomatal pore width to stomatal pore length as described by Wang et al. (2019). All experiments were conducted between 9 AM-1 PM. Results include stomatal aperture values from at least 20 different plants for each time point and treatment (*n* = 500).

### RNA isolation and transcript expression analysis

Four-week-old wild type and *rbohD* plants were subjected to WLS and systemic leaves (leaves number 4, 5, 6 and 7 at ten-leaf rosette stage; Supplementary Figure S3) were collected and immediately frozen in liquid nitrogen at times 0-, 10-, 30- and 60-min. Sixty leaves were pooled from 15 different plants for each biological repeat (three biological repeats were used for each time point and genotype). RNA was extracted using Plant RNeasy kit (Qiagen, Hilden, Germany) according to manufacturer instructions. Total RNA was used for cDNA synthesis (PrimeScript RT Reagent Kit; Takara Bio, Takara Bio, Kusatsu, Japan). Transcript expression was quantified by real-time qPCR using iQ SYBR Green super mix (Bio-Rad Laboratories, Hercules, CA, USA), as previously described (Fichman et al., 2021), with specific primers for the following transcripts: *ZAT10* (AT1G27730), 5′-ACTAGCCACGTTAGCAGTAGC-3′ and 5′-GTTGAAGTTTGACCGGAAGTC-3′; *RAP2.3* (AT3G16770) 5′-AGCAGATCCGTGGTGATAAAG-3′ and 5′-TATACTCCTCCGCCGTCA-3′; *ADH1* (AT1G77120), 5′-GATCATGTGTTGCCGATCTTTAC-3′ and 5′-CTTCTCAGGATCAACACCGAG -3′; *HRE2* (AT2G47520), 5′-GGGAAACGAGAGAGGAAGAATC-3′ and 5′-AAAGGTGTACGTGTCTGGC-3′. *Elongation factor 1 alpha* (5′-GAGCCCAAGTTTTTGAAGA-3′ and 5′-TAAACTGTTCTTCCAAGC TCCA-3′) was used for normalization of relative transcript levels. Results, expressed in relative quantity (2^−ΔΔCT^), were obtained by normalizing relative transcript expression (ΔC_T_) and comparing it to control wild type from local leaf (ΔΔC_T_). The data represent 15 biological repeats and 3 technical repeats for each reaction. SE and Student’s t test were calculated with Microsoft Excel.

### RNA sequencing and data analysis

RNA libraries were prepared using standard Illumina protocols and RNA sequencing was performed using NovaSeq 6000 PE150 by Novogene co. Ltd (https://en.novogene.com/; Sacramento, CA, USA). Quality control for the raw reads was evaluated by FastQC v.0.11.9 (Andrews, 2010) and aggregated with MultiQC tool v.1.13.dev0 (Ewels et al., 2016). Adapter content, ambiguous nucleotide, and any sequences with read length less than 20 bp and a Phred score less than 20 were removed from the raw reads with Trim Galore v.0.6.7 (https://www.bioinformatics.babraham.ac.uk/projects/trim_galore/). The RNA-seq reads were aligned to the reference genome for Arabidopsis thaliana (TAIR10) (downloaded from https://ftp.ensemblgenomes.ebi.ac.uk/pub/plants/release-54/fasta/arabidopsis_thaliana/dna_index/), using HISAT2 short read aligner v.2.2.1 (Kim et al., 2019) which gave a high overall alignment (∼98%). Intermediate file processing of sam to sorted bam conversion was conducted using Samtools v.1.9 (Li et al., 2009). Transcript abundance expressed as fragments per kilobase million (FPKM) was generated using the Cufflinks tool v.2.2.1 from the TUXEDO suite (Trapnell et al., 2012) guided by genome annotation files (downloaded from https://ftp.ensemblgenomes.ebi.ac.uk/pub/plants/release-54/gff3/arabidopsis_thaliana/). Differential gene expression analysis was performed using the Cuffdiff 2 method (Trapnell et al., 2013), also from the same TUXEDO suite. Differentially expressed transcripts were defined as those that had a fold-change with an adjusted *P* ≤ 0.05 (negative binomial Wald test followed by Benjamini–Hochberg correction). For the analysis and visualization of the data, R was used with methods and packages available through CRAN (R Core Team, https://www.R-project.org/) or Bioconductor (Huber et al., 2015). Functional annotation and quantification of overrepresented gene ontology (GO) terms and KEGG pathway enrichment was performed using gprofiler2 package v.0.2.1 (Raudvere et al., 2019) using a threshold of (*P* < 0.05). Venn diagrams were created in Venny 2.1 (BioinfoGP, CNB-CSIC). The different stress-, hormone-, and ROS-response transcripts datasets used for comparisons in Table 1 were obtained from Zandalinas et al., (2019, 2021a), Mustroph et al., 2009 and the GO database using the tool AmiGO (http://amigo.geneontology.org/amigo).

### Statistical analysis

All experiments were repeated at least three times with three biological repeats. Statistical analysis for data presented in Figures 2, 4, 5 and Supplementary Figure S1 was performed using one-way ANOVA followed by Tukey’s post hoc test (*P* < 0.05) in GraphPad. Results are shown as box and whisker plots with borders corresponding to the 25^th^ and 75^th^ percentiles of the data. Different letters denote statistical significance at *P* < 0.05. Statistical analysis for data in Figures 1, 3 and Supplementary Figure S2 was performed by two-sided Student’s t test (\**P* < 0.05) in Microsoft Excel, and results are presented as means ± SE.

## ACKNOWLEDGMENTS

We thank Professor Julia Bailey-Serres for helpful discussion and advice regarding the results. This work was supported by funding from the National Science Foundation (IOS-2110017, IOS-1353886, IOS-1932639), Interdisciplinary Plant Group, and University of Missouri.

## AUTHOR CONTRIBUTIONS

M.A.P.V. and A.T. performed experiments and analyzed the data. M.A.P.V., R.M., T.J., and D.G.M.C. designed experiments, analyzed the data, and/or wrote the manuscript.

## DATA AVAILABILITY

RNA-Seq data was deposited in Gene Expression Omnibus (GEO), under the following accession number: GSE225407.

## SUPPLEMENTARY DATA

**Supplementary Figure S1.**
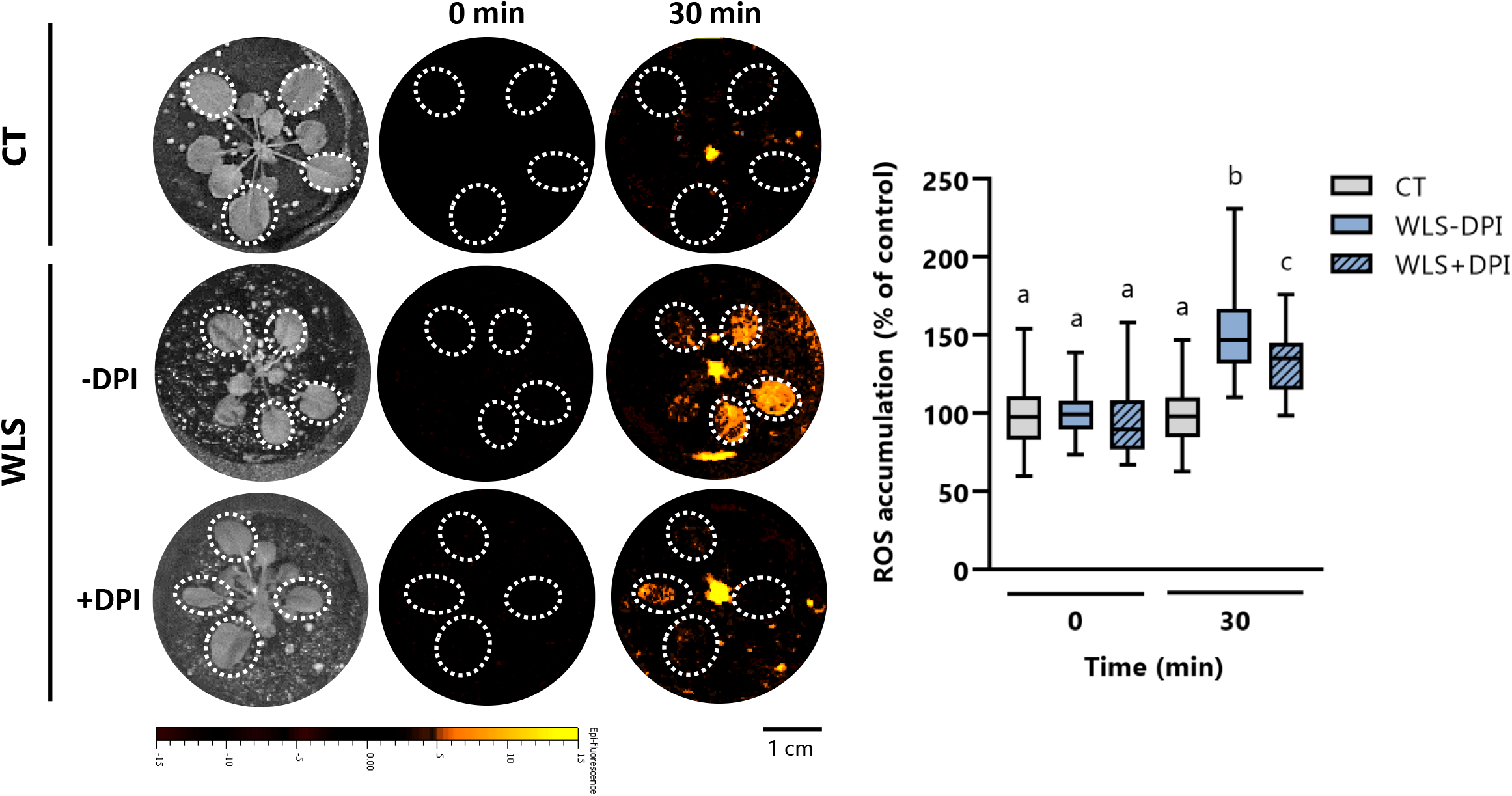
The effect of DPI fumigation on systemic ROS accumulation in response to waterlogging stress (WLS) in wild type plants. Arabidopsis plants were fumigated with a solution containing the dye H_2_DCFDA and 50 µM DPI (+DPI) 30 min before subjecting the plants to WLS. Representative images of ROS accumulation in plants fumigated with or without DPI at 0- and 30-min alongside a box and whisker plots summarizing the results obtain at 0- and 30-min. Borders of box and whisker plots correspond to the 25^th^ and 75^th^ percentiles of the data. Regions of interest are indicated by white ovals in the images. Magnification bar is 1 cm. Different letters denote significance at *P* < 0.05 (ANOVA followed by a Tukey’s post hoc test). All experiments were repeated at least three times with eight plants per repeat. Abbreviations: CT, control; WLS, waterlogging stress; DPI, diphenyleneiodonium; H_2_DCFDA, 2’,7’-dichlorodihydrofluorescein diacetate.

**Supplementary Figure S2.**
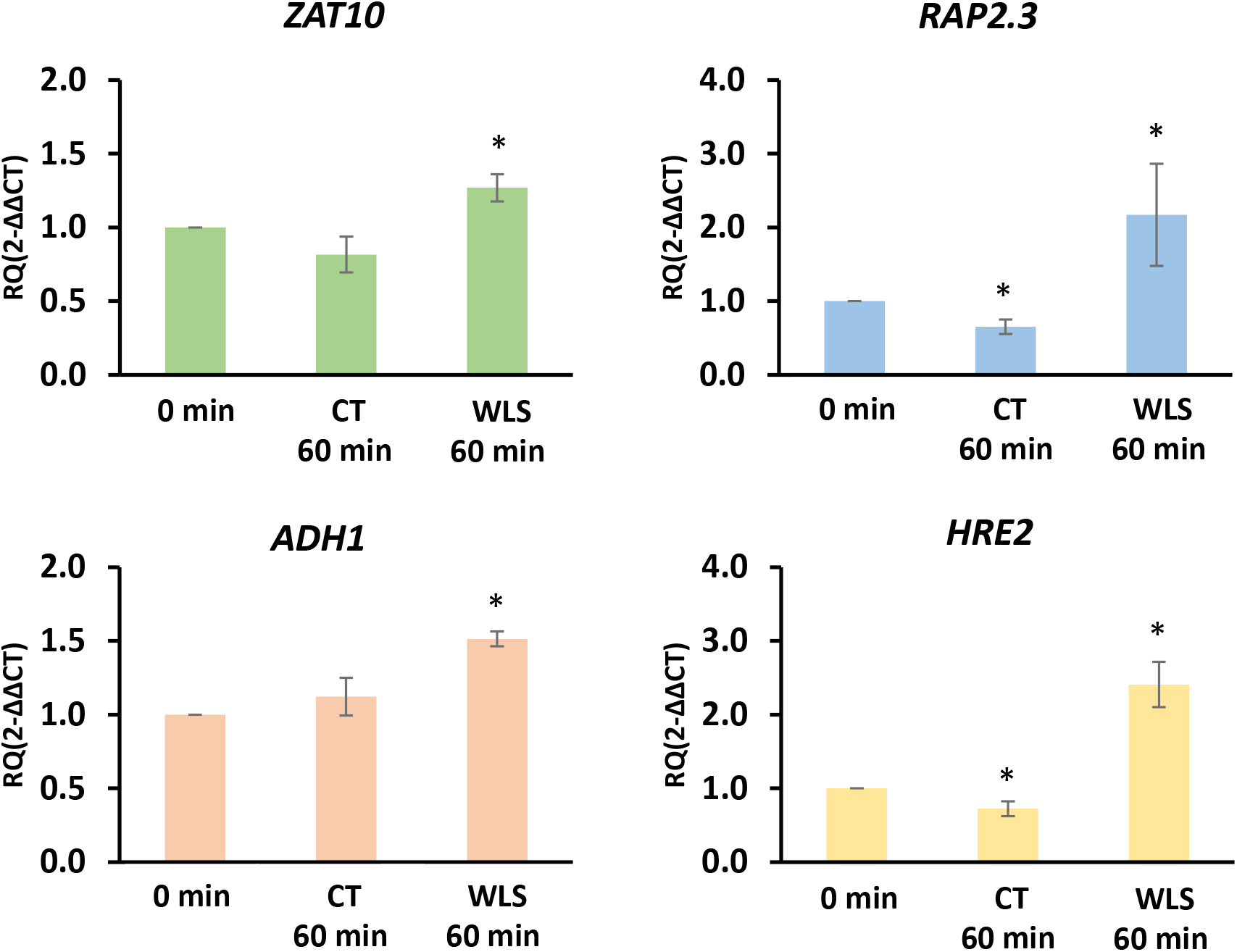
Quantitative reverse transcription polymerase chain reaction (RT-qPCR) analysis of Zinc finger protein 10 (ZAT10), Ethylene response factor subfamily B-2 (RAP2.3), Alcohol dehydrogenase 1 (ADH1), Hypoxia responsive ethylene response factor gene 2 (HRE2), in systemic leaves of wild type plants subjected to waterlogging stress (WLS). Results are expressed as relative quantity (RQ) compared to internal control (elongation factor 1 alpha, EF1α) and time 0. Data represent means ± SE (n = 15, *P < 0.05 according to Student’s t test). CT, control; WLS, waterlogging stress.

**Supplementary Figure S3.**
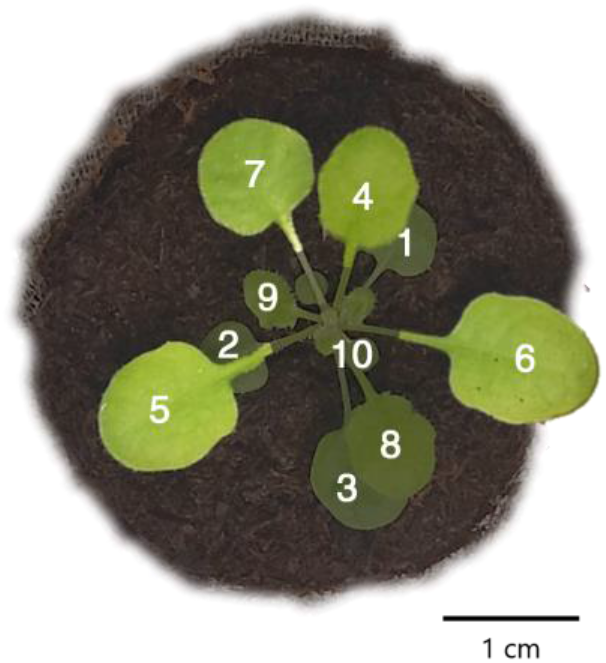
Position of the leaves used for the transcriptomic analysis shown in Figure 5. Systemic leaves number 4, 5, 6 and 7 (highlighted in light green) of control and plants subjected to waterlogging stress (WLS) were collected from wild type and *rbohD* plants and immediately frozen in liquid nitrogen. Magnification bar is 1 cm.

**Supplementary Table S1.** Transcripts differentially expressed in WT plants subjected to waterlogging stress for 10 min (compared to WT 0 min).

**Supplementary Table S2.** Transcripts differentially expressed in WT plants subjected to waterlogging stress for 30 min (compared to WT 0 min).

**Supplementary Table S3.** Transcripts differentially expressed in WT plants subjected to waterlogging stress for 60 min (compared to WT 0 min).

**Supplementary Table S4.** List of transcripts uniquely altered in WT at time 10 min.

**Supplementary Table S5.** List of transcripts uniquely altered in WT at time 30 min.

**Supplementary Table S6.** List of transcripts uniquely altered in WT at time 60 min.

**Supplementary Table S7.** Gene ontology and KEGG pathway enrichment of all the transcripts regulated in WT in response to waterlogging (10 min+30 min+60 min) shown in Fig 6B.

**Supplementary Table S8.** Comparisons between the list of systemic transcripts regulated in WT in response to waterlogging (8,013) with datasets from previous studies with Arabidopsis plants subjected to hypoxia or submergence stress.

**Supplementary Table S9.** Common transcripts found in the overlapping of the transcriptomic response of WT to waterlogging stress with transcripts significantly altered in plants subjected to different stresses, hormone treatments, or ROS (data shown in Table 1).

**Supplementary Table S10.** Transcripts significantly expressed in *rbohD* plants subjected to waterlogging stress for 60 min (compared to rbohD 0 min).

**Supplementary Table S11.** Transcripts found in the comparison between the transcriptional response of WT and *rbohD* plants subjected to waterlogging stress for 60 min (Fig 7A).

**Supplementary Table S12.** Common transcripts found in the overlapping of the transcriptomic response of *rbohD* to waterlogging stress with transcripts significantly altered in plants subjected to different stresses, hormone treatments, or ROS (data shown in Table 1).

**Supplementary Table S13.** Gene ontology and KEGG pathway enrichment of RBOHD-dependent transcripts (1,806+3,028; Fig 7B).

**Supplementary Table S14.** Transcripts altered in their expression at 10, 30 and 60 min and that are RBOHD-dependent (Fig 7C).

